# Offspring overcome poor parenting by being better parents

**DOI:** 10.1101/2022.12.13.520305

**Authors:** Ahva L. Potticary, Christopher B. Cunningham, Allen J. Moore

## Abstract

The evolutionary repercussions of parental effects are often discussed as static effects that can have negative influences on offspring fitness that may even persist across generations. However, individuals are not passive recipients and may mitigate the persistence of parental effects through their behaviour. Here, we tested how the burying beetle *Nicrophorus orbicollis*, a species with complex parental care, responded to poor parenting. We manipulated the duration of parental care received and measured the impact on traits of both F1 and F2 offspring. As expected, reducing parental care negatively affected traits that are ecologically important for burying beetles, including F1 offspring development time and body size. However, F1 parents that received reduced care as larvae spent more time feeding F2 offspring than parents that received full care as larvae. As a result, both the number and mass of F2 offspring were unaffected by the developmental experience of their parents. Our results show that flexible parental care may be able to overcome poor developmental environments and limit negative parental effects to a single generation.

## 1. INTRODUCTION

All parents influence the development of their offspring (Marshall & Uller, 2007, Roach & Wulff, 1987, Mousseau & Fox, 1998, Molinier et al., 2006, Royle et al., 2012), ranging from short-term impacts to effects that persist for multiple generations (Molinier et al., 2006, Cheverud & Moore, 1994, Yin et al., 2019). Adaptive parental effects are expected to evolve to help offspring overcome challenging environments by modifying offspring phenotypes or buffering (Badyaev & Uller, 2009, Mousseau & Fox, 1998, Wilson, 1975). In the former, parents prepare offspring to face a future environment (Potticary & Duckworth, 2020, Mousseau & Dingle, 1991, Dantzer et al., 2013, Burgess & Marshall, 2014), whereas in the latter, parents buffer offspring by compensating for unfavorable conditions that would otherwise be experienced during offspring development. As environments can change rapidly or unpredictably, parental effects on offspring development range from positive (Dantzer et al., 2013, Galloway & Etterson, 2007, Duckworth et al., 2015, Badyaev et al., 2006) to negative (Potticary & Duckworth, 2018, Gluckman et al., 2005, Sheriff & Love, 2013) depending on the environmental context (Bernardo, 1996, Bateson et al., 2014). Consequently, the evolutionary repercussions of parental effects are often viewed as static effects on offspring phenotype (Badyaev & Uller, 2009) that may even persist into subsequent generations. This is an assumption that should be tested (Badyaev & Uller, 2009) because individuals are not passive recipients of their environment. The persistence of a parental effect depends on the interactions between offspring and their environment later in life (Francis et al., 2002, Cheverud & Moore, 1994, Bernardo, 1996), which can also be mediated by flexibility in individual behaviour. For this reason, parental effects that are produced by behaviour, such as parental care, provide a unique opportunity to determine how variation in the behaviour of both parents and offspring influences the persistence of parental effects. Here, we specifically investigate how behaviour may impact the persistence of parental effects over time.

Parental care is expected to buffer offspring development from environmental threats (Wilson, 1975). Moreover, parental care by definition influences offspring development, and variation in parental care can have cascading effects that impact multiple generations (Champagne, 2008, Rossiter, 1996). The goals of parents and offspring are often not aligned, as variation in parental care can be driven by environmental conditions that change the benefits of allocating effort to current and future reproductive opportunities (Stearns, 1989, Richardson et al., 2020). Thus, on one hand, parental care is a coordinated strategy to buffer offspring from current threats, and on the other, variation in parental care can induce transgenerational effects that may even be negative, depending on the ecological context. This is particularly the case because parental care is not a single behaviour but a composite of multiple behaviours that may buffer or modify offspring development in different ways. Despite the long-standing hypothesis that parental care acts as a buffer (Wilson, 1975, Tallamy, 1984), it is unclear how parental care may buffer offspring development within and between generations. One way to test these ideas is to manipulate the parental care that young receive and determine how this impacts the expression of parental care and F2 offspring phenotypes when these individuals are parents themselves.

Here, we investigated how variation in parental care influences the development and behaviour of the burying beetle *Nicrophorus orbicollis* across generations. Specifically, we tested whether individuals that received poor parenting can compensate for their own developmental environment when they are parents or whether the developmental impact of poor parenting is perpetuated to the next generation. Given that parental care is a complex behavioural trait, we further asked how the expression of the behaviours composing parental care are affected by an individual’s developmental experience. We investigated this by manipulating the duration of parental care that mixed foster broods received to produce larvae that experienced reduced (six hours) or full (approximately seven days) parental care. Burying beetles require small animal carcasses to breed and provide complex care to their young, which includes a suite of parenting behaviours (Robertson, 1993, Eggert & Müller, 1997, Scott, 1998). In *Nicrophorus*, parents bury and defend carcasses, which in part acts to ensure that the carcass is available for longer (Robertson, 1993, Trumbo & Sikes, 2021), eliminate bacterial and fungal consumers to avoid putrefaction (Haberer et al., 2014, Arce et al., 2012), create cavities with predigested carrion “soup” to support offspring self-feeding (Duarte et al., 2021, Capodeanu-Nägler et al., 2018, Eggert & Müller, 1997), and directly provision food to larvae through regurgitation of carrion (Eggert et al., 1998). Both direct provisioning and maintenance of the carcass are known to increase larval size and growth rate in *Nicrophorus* (Lock, 2012, Lock et al., 2004, Walling et al., 2008). Parental behaviour affects not only larval development but also has a significant impact on the traits of those offspring when they become adults. Larvae do not feed after they leave the carcass until they metamorphose into adults; thus, adult body size is largely determined by resource acquisition during the larval period, similar to other insects (Nijhout, 2003). Body size is an important trait for *Nicrophorus*, as larger beetles are more effective competitors for carcasses, which are ephemeral and are expected to be a limited breeding resource (Otronen, 1988, Robertson, 1993, Royle & Hopwood, 2017). Thus, parental care is particularly important in this system because it creates a direct link between a larva’s development and competitive ability upon reaching adulthood.

Our study was designed to determine whether poor parenting begets poor parenting or if parents can behaviourally buffer their offspring from their own developmental experience. Both hypotheses predict that removing parental care will impact ecologically relevant F1 offspring traits. If poor parenting in one generation leads to poor parenting in the next, then parents that receive reduced parental care should show less care than individuals raised in full care environments, and this variation in care should produce concomitant changes in F2 offspring phenotypes. Alternatively, if parents can behaviourally buffer their offspring, then parents that received reduced care should differ in their expression of care behaviours relative to full care parents, and there should be no difference in F2 larval traits between treatments. As expected, we found that our experimental reduction of parental care greatly reduced the body size of F1 offspring when they became adults. However, we also found that when these individuals became parents, they altered the expression of key parental behaviours and, consistent with buffering, raised broods of equal number and weight. These results suggest that parental care produces an adaptive parental effect that ensures that the negative effects of poor parenting do not persist. Thus, parental care may not only buffer offspring development within a generation but may also act as a buffer that mitigates the legacy of environments past.

## 2. MATERIALS AND METHODS

### 2.1 Insect colony and husbandry

We used *Nicrophorus orbicollis* from an outbred colony maintained at the University of Georgia, which is regularly supplemented with wild-caught beetles from Whitehall Forest in Athens, GA, USA. All beetles were maintained in an incubator (Percival, model no. I4-66VL, Perry, IA) at 22 ± 0.1°C on a 14:10 hours light:dark cycle that simulated summer breeding conditions. We bred all individuals used here in the laboratory to ensure that parentage and age were known and to standardize rearing conditions. We housed beetles individually at larval dispersal in 9-cm-diameter by 4-cm-deep circular compostable deli containers (EP-RDP5, Eco-Products) filled with approximately two cm of organic potting soil. The larvae had no further social interactions with other burying beetles until they were randomly allocated into treatments. We defined individuals as day 1 of adulthood on the day that they eclosed. Beetles were fed *ad libitum* with organic ground beef twice a week following adult eclosion, i.e., the emergence of the adult beetle.

### 2.2 Effect of removing parental care on offspring development

We manipulated the duration of parental care to determine the effect of parental care on F1 offspring traits. To do this, we set up 36 families, N = 18 in both the full and reduced care treatments. We randomly mated focal virgins on days 13-19 of adulthood by placing an unrelated male and female in a small plastic container and observing them to ensure that mating occurred. The pair were then moved to a plastic breeding box filled with approximately two cm of potting soil moistened with distilled water and containing a freshly thawed mouse 20.8 ± 0.6 g SD (max = 22.2, min = 20.0; Rodent Pro, Inglefield, IN, U.S.A.). There was no difference in the size of the mice provided between treatments (20.9 g in the full care treatment, 20.8 g in the reduced care treatment; *F*_1,34_ = 0.153, P = 0.698). Breeding boxes were then returned to the incubator and kept on a darkened shelf to simulate underground breeding conditions. We checked the breeding boxes for eggs every twenty-four hours. Once we observed eggs, we moved the female and brood ball to a new breeding box and removed the male. We left the eggs in the original breeding box with a small piece of ground beef. After 90 hours had elapsed since egg laying, we checked the breeding box containing the eggs every hour to determine when the larvae had hatched and crawled to the beef. After we observed that the focal females’ larvae had hatched, we provided the focal female with an unrelated, mixed brood of 10 larvae. Burying beetles identify their offspring temporally and eat any larvae that arrive too early on the carcass (Oldekop et al., 2007, Potticary et al., 2023); thus, we only provided females with larvae after her own larvae had hatched. For the full care treatment, we allowed females to associate with larvae until at least two of the larvae had dispersed from the breeding carcass (∼7-8 days). For the reduced care treatment, we allowed females to associate with larvae for 6 hours and then removed the females. There is significant variation in the duration of care that parents provide in the wild (Eggert & Müller, 1997, Wilson & Fudge, 1984), and parents can desert early in the post-hatching stage. We checked the experimental treatments daily to determine the larval dispersal date and larval survival.

We measured several ecologically relevant offspring traits (Lock et al., 2004), including duration of developmental periods, mass at dispersal, and adult size. We checked the treatments daily to determine the number of surviving larvae and to measure the duration of each developmental stage. The development of burying beetles occurs in discrete and easily scored periods: from hatching to dispersal from the carcass, during which time the larvae are fed by the parents and feed from the carcass (*hatch to dispersal*); non-feeding “wandering” larvae that have dispersed from the carcass and then bury themselves at a distance from the consumed carcass where they form pupae (*dispersal to pupae*); and pupal development to adult eclosion, where they emerge as winged teneral adults from the soil (*pupae to eclosion*). After the larvae had dispersed from the carcass, we measured larval mass to the nearest 0.1 mg and placed larvae into individual containers. We determined the sex and measured the pronotum length of full and reduced care offspring on days 14 – 16 post-eclosion with calipers to the nearest 0.01 mm.

### 2.3 Effect of variation in parental care environment on the next generation

To determine the effect of the experimental manipulation of parental care on F1 offspring when they reached adulthood, we established 89 uniparental broods using the resulting offspring from the reduced and full care treatments (26 full care males, 20 full care females, 24 reduced care males, and 17 reduced care females). Individuals were randomly selected from the families within each treatment. On days 14 – 16 following eclosion, adult beetles that received full or reduced care as offspring were provided with an unrelated, virgin mate from our stock colony and a 19.6 - 21.2 g thawed mouse carcass (mean = 20.6 g). These were placed in a breeding container with approximately two cm of moistened soil. As variation in parental care may manifest as variation in investment across the breeding cycle, we fitted breeding boxes with an escape chamber to allow focal beetles to desert their breeding attempt. We checked breeding boxes daily for eggs. We placed beetles that escaped into the escape chamber before egg laying back into the breeding box, while we counted beetles that escaped after egg-laying and remained in the escape chamber for more than 15 minutes as deserters and removed them. Once we observed larvae, we removed the non-focal partner, which left the focal male or female in a uniparental state to care for the offspring. If the focal beetle deserted a breeding attempt before larvae hatching, we did not conduct any no further monitoring on that breeding attempt.

We measured the expression of parental care using behavioural observations twenty-four hours after the larvae were first observed on the carcass. We moved the breeding boxes into an observation room and allowed the focal beetle to acclimate for at least 30 minutes before observation. All observations were conducted under red lights to ensure no light effects on individual care behaviour, as *N. orbicollis* is nocturnal (Wilson et al., 1984). In each behavioural observation, we recorded both direct and indirect parental care behaviours for 15 minutes (Walling et al., 2008). We defined direct care as when the parent fed regurgitated carrion to the larvae. We defined indirect care as activities related to carcass cleaning and maintenance, which allows for the preservation of the carcass for future offspring provisioning and thus represents a type of parental care without direct interactions with offspring. We also recorded the time spent showing “no care,” defined as when the parent was on the carcass but not engaged in direct or indirect care, or when the individual was off the carcass. We recorded behaviours continuously as both a duration and frequency (e.g., the number of times an adult fed larvae). This gave each individual three scores (direct care, indirect care, and no care) which together summed to 15 minutes.

We did not manipulate the number of offspring in the F2 generation. To account for the fact that rates of parental provisioning may be affected by the number of larvae (Rauter & Moore, 2004), we also measured the amount of time larvae spent on the carcass as well as the number and mass of larvae at dispersal for each larvae as a proxy for parental effort. Measurements of development times and larval mass were the same as those used in the experimental manipulation of the duration of parental care (see the previous section).

### 2.4 Statistical analyses

We performed all statistical analyses using JMP Pro (v. 16.0.0, http://jmp.com) and produced figures in SigmaPlot (v. 14.5, http://www.sigmaplot.co.uk). Results are presented as the means ± standard error (SEM) unless otherwise noted. All analysis of variance (ANOVA) tests were two-tailed.

#### 2.4.1 Effect of removing parental care on offspring development

We used nested ANOVA to analyze the effect of care treatment (reduced or full parental care) on F1 larval mass, with care treatment and family as random factors. We used family as a nested term to control for measures of multiple individuals in each family. We analyzed the duration of the *hatch to dispersal* period as a single value for each family, as all larvae disperse from the carcass within hours of each other. We analyzed the duration of *hatch to dispersal* with a Pearson chi-square goodness-of-fit test, as these data were categorical (see Results). The other periods of development can vary between individuals within a family (Lock, 2012); therefore, we analyzed each larva individually using nested ANOVA with family nested under treatment. We included larval mass at dispersal as a random factor when analyzing development off the carcass and total postembryonic development, including *dispersal to pupae*, *pupae to eclosion*, and *hatch to eclosion*, as the duration of post-dispersal development can be influenced by mass (Rauter & Moore, 2002). The relationship between an individual’s mass at larval dispersal and their body size in adulthood was analyzed using linear regression.

#### 2.4.2 Effect of variation in parental care environment on the next generation

We tested the hypothesis that the parental care treatment (reduced or full care) received by F1 beetles influences their behaviour when they become parents and the development of their offspring (F2 generation). We analyzed whether F1 parents raised with reduced or full care differed in their abandonment of larvae using a chi-square test. We then analyzed the effect of F1 care treatment on the behaviour of F1 towards F2 offspring using a two-way ANOVA with the sex of the F1 parent and care treatment as factors, with the proportion of the behavioural observation that the parent spent a) on the carcass (total time on the carcass, regardless of behaviour expressed), b) engaged in indirect care, or c) engaged in direct care behaviour, as the dependent variable.

We tested the hypothesis that the parental care treatment received by F1 beetles influences the development of their F2 offspring in several ways. First, we analyzed brood size and F2 larval mass with a fully factorial two-way ANOVA with caregiver sex as a fixed factor and care treatment as a random factor. We used the average F2 larval mass for the family reared by each F1 parent to control for the fact that multiple individuals were measured in each family. We analyzed the duration of F2 larval development on the carcass with F1 caregiver sex, care treatment, and their interaction using a GLM with a Poisson distribution, as these data are categorical because larvae within a family disperse on the same day.

We next tested how F1 behaviour to their F2 offspring influenced the number of surviving larvae and their mass at dispersal. Average larval mass and the number of offspring are strongly correlated (see Results) because the rates of provisioning can be influenced by the number of begging offspring and impact offspring growth (Lock, 2012, Lock et al., 2004, Smiseth et al., 2003). For this reason, we used a principal component analysis to reduce them to a single variable. PC1 was retained because it had an eigenvalue greater than one and explained 84% of the variance. We used Pearson correlation to determine whether direct and indirect care behaviours were related to PC1.

## 3. RESULTS

### 3.1 Effect of reducing parental care on offspring development

Larval survival on the carcass was high for both the full and reduced care treatments; from an initial ten fostered larvae placed on the carcass, three to ten larvae survived to dispersal from the mouse with a mean of 8.3 ± 0.4 surviving larvae for full care families and 8.7 ± 0.4 for reduced care (*F*_1,34_ = 0.430, P = 0.516). Following dispersal, 12% (N = 35 of 304) of the total individuals died before adult eclosion, including 11% of larvae that received full care (N = 16 of 148) and 12% of larvae that received reduced care (N = 19 of 156).

The care treatment F1 larvae received - reduced or full care - affected both the duration of development on the carcass and larval mass at dispersal, but the duration of developmental periods that occurred once larvae had left the carcass was primarily influenced by larval mass. F1 larvae that received full care completed development on the carcass faster than larvae that received reduced care (*hatch to dispersal*; χ^2^ = 18.857, df = 2, p < 0.001, N = 36 families). Larvae in the full care treatment dispersed in 7 or 8 days, whereas reduced care larvae dispersed in 8 or 9 days. F1 larval mass at dispersal was strongly and positively affected by the duration of parental care received. Controlling for the significant effect of variation among families (*F*_34,269_ = 10.155, P < 0.0001), larvae that received full care were larger than those that received reduced care (*F*_1,34_ = 45.358, P < 0.0001; **Figure 1*A***). Controlling for variation amongst families, larvae that received full care weighed an average of 500.4 ± 4.4 mg, whilst larvae that received reduced care weighed an average of 373.4 ± 4.2 mg. F1 larval mass at dispersal was positively and strongly correlated with their final adult body size (*r* = 0.91, N = 248).

**Figure 1.**
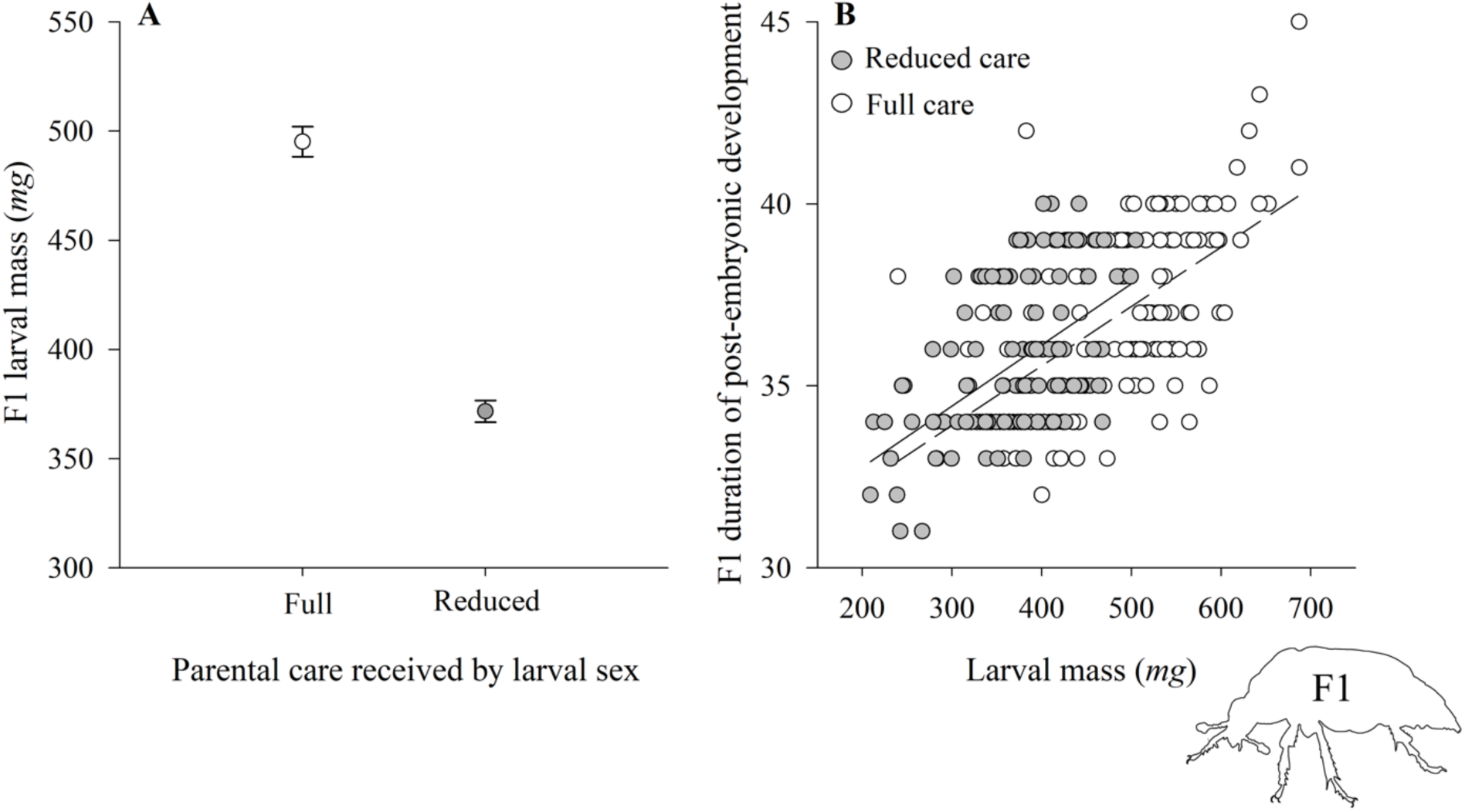
Care influences larval development time and size. Data from F1 beetles that experience full or reduced parental care; generation indicated in beetle graphic. ***(A)*** Larvae that received full care were larger than larvae that received reduced care. Error bars are mean ± SEM. *(**B)*** Duration of the larval period (in days) from larval hatch until ecolsion was statistically dependent on larval mass at dispersal but not on care treatment (N = 248). Grey circles and the solid line refer to larvae that received reduced care, while open circles and the dashed line refer to individuals that received full care.

The time that larvae spent in the *dispersal to pupation* stage showed statistically significant variation attributable to family (*F*_34,261_ = 9.301, P < 0.0001) and larval mass at dispersal (*F*_1,261_ = 34.501, P < 0.0001), but not to care treatment (*F*_1,261_ = 3.363, P = 0.073). The same was true for the duration of the *pupae to adult eclosion* stage, with statistically significant variation arising from family (*F*_34,231_ = 3.173, P < 0.0001) and larval mass at dispersal (*F*_1,231_ = 55.586, P < 0.0001), but not to care treatment (*F*_1,34_ = 0.488, P = 0.487). Even though larvae in the full care treatment developed faster on the carcass, the duration of total postembryonic development, *hatch to eclosion*, was influenced primarily by family (*F*_34,232_ = 9.699, P < 0.0001) and larval mass at dispersal (*F*_1,232_ = 77.283, P < 0.0001), and not to care treatment (*F*_1,34_ = 1.311, P = 0.254; **Figure 1*B***).

### 3.2 Effect of variation in parental care environment on the next generation

There is no evidence that manipulation of care experiment (full or reduced care) influences whether parents abandoned the breeding attempt before showing parental care to the larvae. Focal male caretakers were more likely to abandon before eggs hatched (reduced care males: 11/24 breeding attempts; full care males: 11/26 breeding attempts) than female caretakers (reduced care females: 2/17 breeding attempts; full care females 2/20 breeding attempts). Once the larvae hatched, most of the focal parents stayed for the entire larval development period on the carcass; when caretakers did abandon larvae, they were almost exclusively males (10/29 males, 1/33 females; χ^2^ = 11.640, df = 1, P = 0.0012). The one female that abandoned early had received full care; for males, four had received full care and six received reduced care.

There are strong differences between uniparental females and males in this species (**Figure 2**) and F1 care treatment affected some parental care behaviours. Females spent more time on the carcass during the observation period than males (*F*_1,55_ = 17.610, P < 0.001; **Figure 2*A***), regardless of the care treatment they received as larvae (*F*_1,55_ = 1.878, P = 0.176), and there was not a statistically significant interaction between care treatment and caretaker sex on the amount of time spent on the carcass (*F*_1,55_ = 1.332, P = 0.253). The time that parents spent regurgitating to their larvae, or direct care, was influenced by F1 sex and care treatment, where caretakers that received reduced care provided more direct care themselves. Females provided more direct care than males under both treatments (*F*_1,55_ = 12.699, P = 0.0008; **Figure 2*B***), and direct care was strongly influenced by care treatment (*F*_1,55_ = 6.761, P = 0.012). This pattern was consistent for both males and females, but there was no interaction between caretaker sex and care treatment (*F*_1,55_ = 0.111, P = 0.741). In contrast, indirect care, or time spent maintaining the carcass, was not affected by caretaker sex (*F*_1,55_ = 3.072, P = 0.085; **Figure 2*C***) or care treatment (*F*_1,55_ = 1.181, P = 0.282), nor was there an interaction between sex and care treatment (*F*_1,55_ = 0.705, P = 0.405).

**Figure 2.**
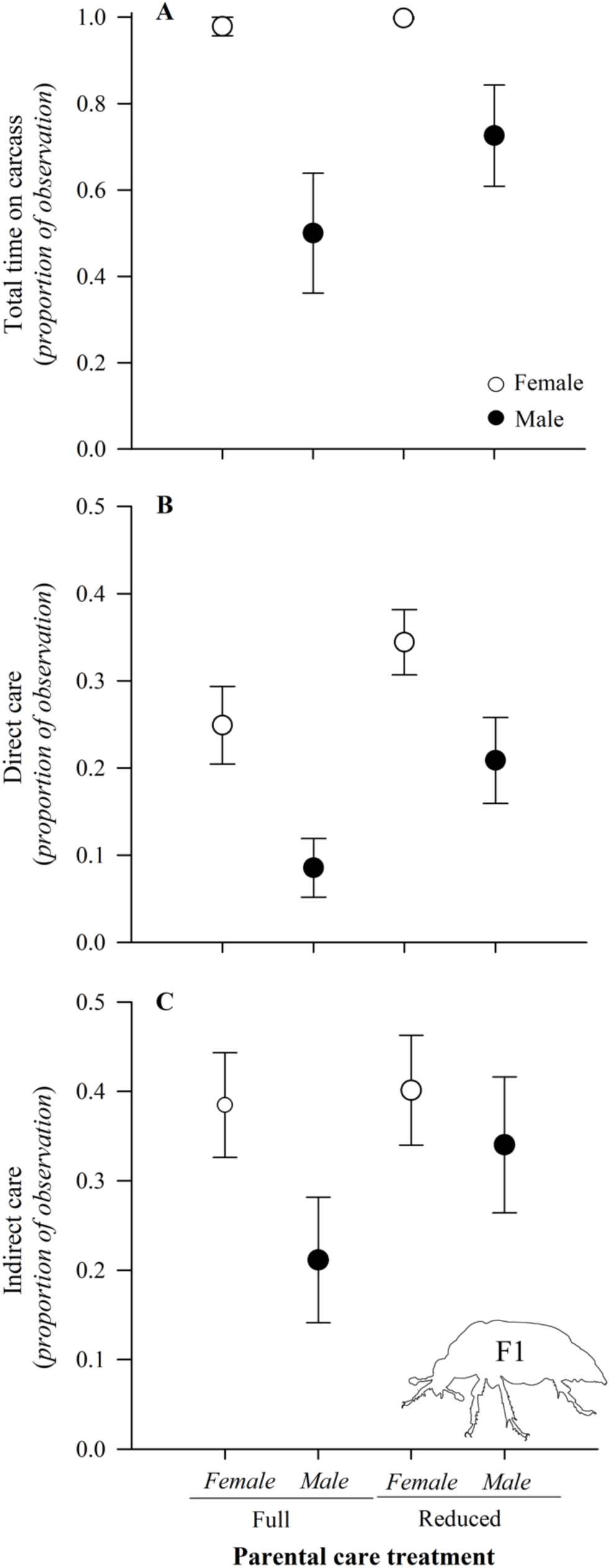
Parental behaviour of F1 parents is affected by the care received as larvae. Data from F1 beetles that experience full or reduced parental care; generation indicated in beetle graphic. N = 15 observations for all treatments except full care males, where N = 14 observations. ***(A)*** Females spent more time on the carcass during the observation period than males regardless of how much care they received. In addition, ***(B)*** both females and males that experienced reduced care increased the amount of direct care given to their offspring but ***(C)*** not indirect care. Error bars are mean ± SEM.

The effects of the reduced and full care treatments experienced by the F1 generation did not carry over into the F2 generation. There was no statistically significant effect on the duration of the *hatch to dispersal* period arising from F1 caregiver sex (χ^2^ = 0.405, df = 1, P = 0.524), F1 care treatment (χ^2^ = 0.004, P = 0.949), or their interaction (χ^2^ = 0.405, P = 0.524). There were no significant effects on the average mass of dispersing F2 larvae arising from F1 caregiver sex (*F*_1,57_ = 4.313, P = 0.286), F1 care treatment (*F*_1,57_ = 2.728, P = 0.347), or their interaction (*F*_1,57_ = 0.513, P = 0.477; **Figure 3*A***). Female caregivers reared more larvae to dispersal than males (*F*_1,57_ = 19.527, P < 0.0001), but there was no effect of F1 care treatment (*F*_1,57_ = 0.567, P = 0.455), or their interaction (*F*_1,57_ = 0.147, P = 0.703; **Figure 3*B***).

**Figure 3.**
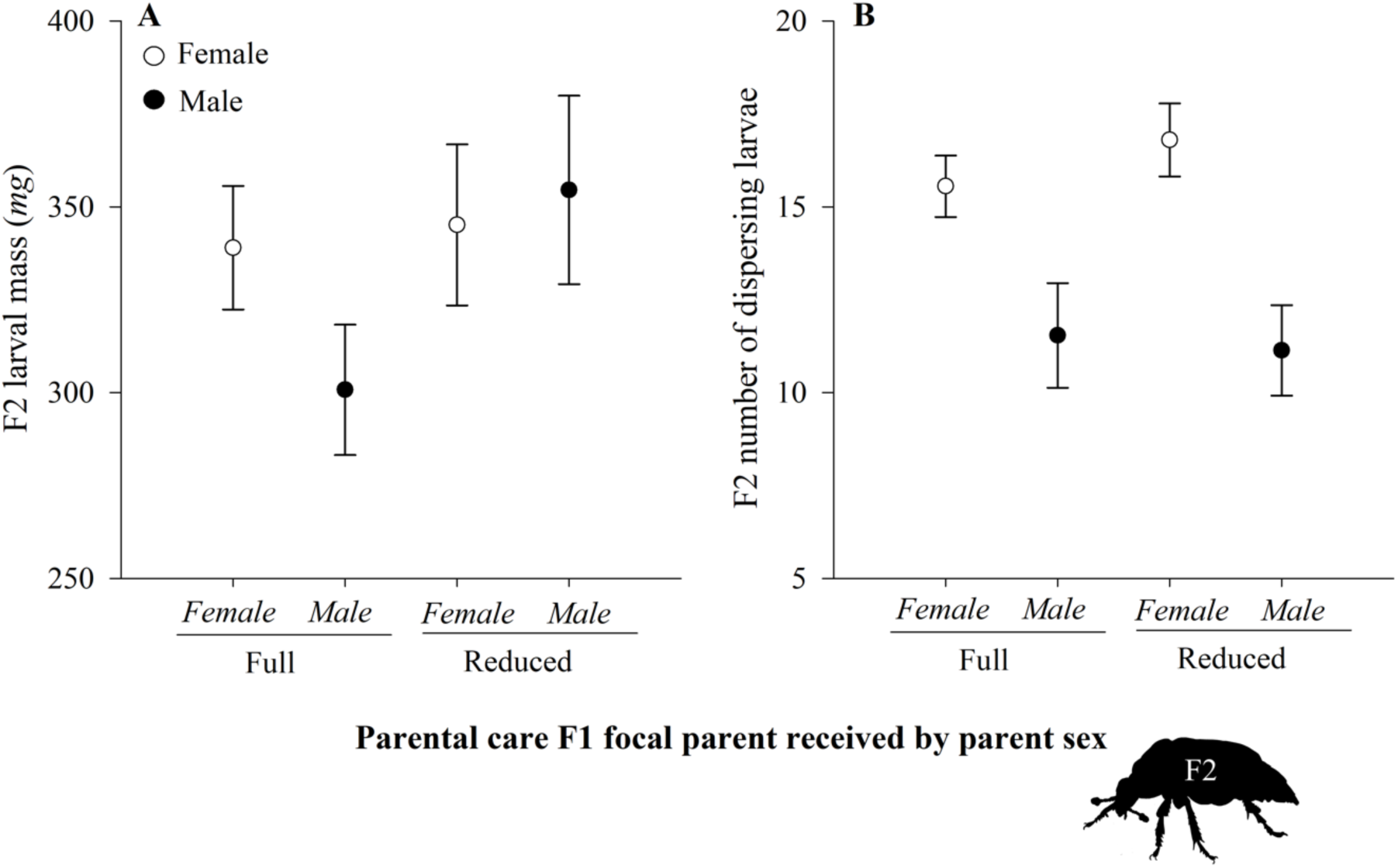
Parent sex influences offspring outcomes, but not the parental developmental environment. Data are from larvae in the F2 generation, whose F1 parents had experienced full or reduced parental care; generation is indicated in the beetle graphic. **(*A*)** There was no difference in the average larval mass of F2 offspring, regardless of whether the F1 parent had received full or reduced care. (***B)*** Females reared more offspring than males, but there was no effect of F1 care treatment on the number of F2 offspring reared. Female parents (N = 18 full care, N = 15 reduced care) are indicated with open circles and male parents (N = 13 full care, N = 15 reduced care) with closed circles. Error bars are mean ± SEM.

The number of larvae that survived to dispersal was strongly and negatively associated with average larval mass at dispersal (linear regression; *r*^2^ = 0.47, P < 0.0001, n = 60). For this reason, we created a composite variable, PC1, to reflect the tradeoff between the number of larvae and their mass. We found that parental care behaviours were strongly associated with offspring outcomes. There was a negative relationship between the amount of direct care F1 parents showed to F2 larvae and PC1 (*r* = −0.55, P < 0.0001; **Figure 4**), indicating that parents that showed more direct care had more larvae survive to dispersal, but parents with more larvae typically had relatively smaller larvae. There was no relationship between the indirect care that F1 parents showed to their larvae and PC1 (*r* = −0.05, P = 0.73), indicating that there was no relationship between the amount of time parents spent maintaining the carcass, larval mass, and the number of larvae to disperse.

**Figure 4.**
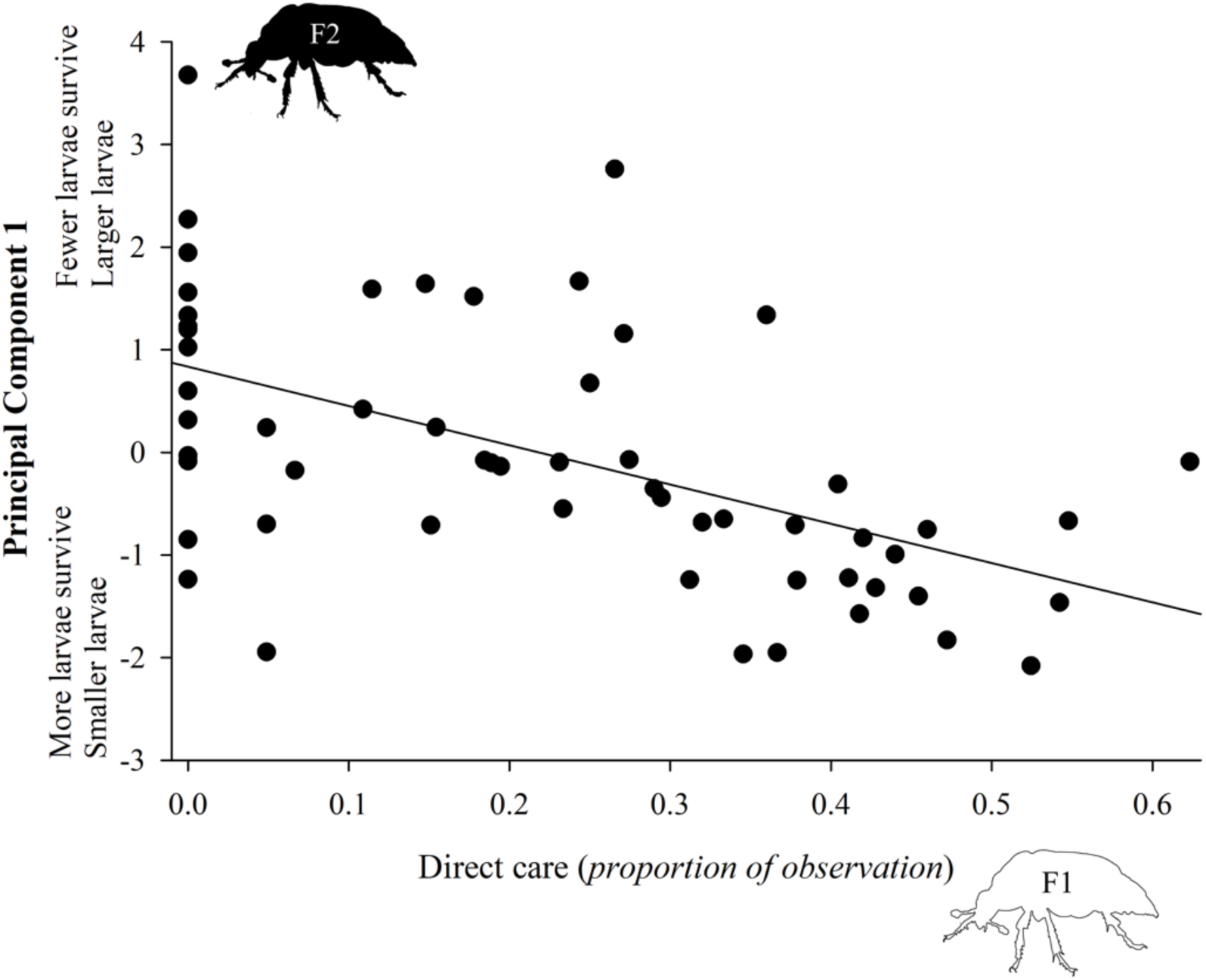
Parents that showed more direct care had more, but relatively smaller, larvae survive to dispersal. Data for F1 beetles expressing direct care toward F2 larvae; generation indicated in beetle graphics on each axis.

## 4. DISCUSSION

Adaptive parental effects are expected to evolve to help offspring overcome challenging environments by modifying offspring phenotypes or buffering (Mousseau & Fox, 1998, Wilson, 1975, Badyaev & Uller, 2009, West-Eberhard, 2003). We tested whether poor parenting begets poor parenting or if parents can compensate for their own poor developmental environment when they become parents themselves by changing their parental behaviour. Our manipulation of the parental care environment influenced the development of ecologically relevant offspring traits, including the duration of development on the carcass and body size (**Figure 1**). However, there was no difference in the size or number of offspring in the next generation when the affected individuals became parents themselves (**Figure 3**). Parents that received reduced parental care spent more time directly transferring nutrients to offspring, as opposed to parents that received full care, who invested relatively more in indirect mechanisms to enable nutrient acquisition by the larvae (i.e., maintenance of the food resource; **Figure 2**). Moreover, parents that spent more time in direct care had more larvae survive to dispersal (**Figure 4**), which is concordant with previous results demonstrating that direct provisioning increases offspring size or growth rate in *Nicrophorus* species (Lock, 2012, Lock et al., 2004). Together, these data suggest that parents may behaviourally compensate for their own poor developmental environment, an adaptive parental effect that may ensure that poor developmental environments do not persist.

Beetles that received full parental care were larger and developed faster on the carcass, although larger beetles overall took longer to develop into adult beetles. Why might the duration of development on the carcass differ from other developmental periods? On a local level, developmental rates can be directly influenced by variation in parental care. Parents regurgitate to larvae, and direct provisioning has been shown to increase offspring size or growth rate across taxa (Naef-Daenzer & Keller, 1999, Lock, 2012, Lock et al., 2004), and we similarly found a pattern between the time invested in direct care and the number of surviving larvae (**Figure 4**). An ultimate explanation for rapid larval development on the carcass may be that there is selection arising from competition or predation for offspring to leave the carcass quickly. Studies of *N. orbicollis* in the wild find that nests are frequently discovered by scavengers or taken over by conspecifics, which generally eat larvae and eggs during successful takeovers (Robertson, 1993). High predation risk has been associated with rapid development rates in nests across bird species (Martin et al., 2011). Thus, we suggest that local provisioning rates serve to enhance offspring growth rates while on the carcass, reducing the risk at this vulnerable stage while still allowing larvae to attain large adult size.

Our experimental reduction of parental care resulted in F1 larvae that took longer to develop on the carcass, were smaller as adults, and invested more time in direct care of offspring than beetles that received full parental care. However, despite an observed increase in direct provisioning by reduced care beetles, there was no difference in the mass or number of F2 offspring between reduced and full care parents. Why did the increase in direct provisioning by reduced care parents not result in larger offspring than full care parents, which provisioned less? One possibility is that behaviour is not the causative factor influencing offspring development, although this explanation is not parsimonious given that a similar experiment in *N. vespilloides* found that reducing the parental care an individual received resulted in poor parenting in later life (Kilner et al., 2015), removing parental care altered both development on the carcass and body size (this study), and body size is associated with the extent of parental care across *Nicrophorus* (Jarrett et al., 2017). Alternatively, the increase in direct provisioning may allow parents to compensate for their developmental environment, particularly if receiving reduced parental care impacts other traits, such as egg size. Steiger (2013) found that smaller *N. vespilloides* females lay smaller eggs than larger females. A tradeoff between allocation to eggs and to post-hatching care may explain why reduced care parents worked harder than full care parents and yet had similar numbers and sizes of offspring survive to dispersal.

Parental effects range from short-term effects that dissipate when offspring pass a certain developmental stage (Gluckman et al., 2005) to effects that persist for multiple generations, even for the same traits (e.g., stress reactivity; Meaney, 2001, Mustoe et al., 2014, Sheriff et al., 2017). Previous studies on other *Nicrophorus* species and taxa have documented such transgenerational effects of poor parental care environments (e.g., Lock, 2012, Kilner et al., 2015, Champagne, 2008, Meaney, 2001). An unanswered question is why the effects of a poor developmental environment are propagated across generations under some circumstances but not in others. The persistence of parental effects likely depends on both abiotic and biotic factors, such as the durability of the physiological mechanisms that transmit them (Weaver et al., 2004, Horsthemke, 2018) and concordance between the parent and offspring environments (Bateson et al., 2014, Marshall & Uller, 2007, Gluckman et al., 2005). Moreover, the persistence of parental effects likely relies on whether the effects arise from an evolved interaction between parents and offspring development or are merely a disruption of normal development (Gluckman et al., 2005). Adaptive parental effects are predicted to induce variation in offspring traits that are functionally related to the environment that triggers them (e.g., Sharda et al., 2021, Duckworth et al., 2015, Tollrian, 1995, Sheriff et al., 2017). In our study, *N. orbicollis* that received poor parenting altered their parental feeding behaviour, and there were no differences between the F2 offspring of parents raised with reduced or full care. These data imply that parental care may act as a developmental buffer between generations and indicate evolution for this purpose in *N. orbicollis,* particularly given how important body size is for competitive interactions in burying beetles (Robertson, 1993, Creighton, 2005, Schrempf et al., 2021). While many processes may alter the persistence of parental effects, changes in the expression of parental care may enable the reversal of poor developmental environments.

